# Rationally designed minimized TbpB confers broad protection against meningococcal infection

**DOI:** 10.1101/2025.10.30.685603

**Authors:** Epshita A. Islam, Jamie E. Fegan, Gregory B. Cole, Charles Calmettes, Dixon Ng, Natalie Y. T. Au, Carolyn M. Buckwalter, Sang K. Ahn, David M. Curran, Laura-lee Caruso, Anthony B. Schryvers, Trevor F. Moraes, Scott D. Gray-Owen

## Abstract

Transferrin binding protein B (TbpB), an iron acquisition protein, has long been recognized as a promising vaccine candidate targeting the pathogenic Neisseria species, including *Neisseria meningitidis*, the cause of meningococcal disease, and *Neisseria gonorrhoeae*, the cause of gonorrhea. A challenge to the development of this protein as a vaccine immunogen is the extent of antigenic variability it exhibits, which complicates the selection of a single variant to elicit a broadly cross-protective immune response. We have utilized structure-informed antigen engineering to develop a minimized version of TbpB consisting of the protein’s carboxy-terminal lobe with its variable surface loops removed. Here, we reveal the effectiveness of this “loopless C-lobe” as an independent immunogen, with structural characterization and stability studies to demonstrate its integrity, and murine immunization and challenge studies that establish its ability to elicit robust protective efficacy by using *N. meningitidis* invasive infection and nasopharyngeal colonization models. The breadth of protection provided, as measured by both in vitro analysis and cross-protection mouse challenge studies, indicate that a single loopless C-lobe elicits a broadly cross-protective immune response against the diverse panel of meningococcal strains tested, and that the cross-reactivity is superior to that offered by the intact TbpB or the native C-lobe. Together, this study demonstrates the utility of structure-informed antigen engineering towards the development of broadly efficacious protein-based vaccines.

**Importance:** Surface-exposed proteins on bacterial pathogens are enticing candidate vaccine targets, however their exposure to the immune system frequently leads to high levels of antigenic variation, a factor that complicates the development of broadly protective vaccines. Here, we undertake an antigen engineering approach to develop a minimized version of a surface lipoprotein, transferrin binding protein B, where variable regions of the protein have been removed to focus the immune response to conserved regions of this antigen. We combine structural studies and mouse infection models of *Neisseria meningitidis*, the cause of meningococcal disease, and *Neisseria gonorrhoeae*, the causative agent of gonorrhea, to reveal that our strategic minimizing of the protein immunogen focuses the immune response to extend the resulting breadth of cross-reactivity and cross-protection.

## Introduction

Surface lipoproteins (SLPs) have long been viewed as promising vaccine targets for gram-negative bacterial pathogens, including the pathogenic *Neisseria* species, due to their surface accessibility, importance in pathogenesis, and inherent stability as potential immunogens (Lissolo et al., 1995; Rokbi et al., 1997; Steere et al., 1998; Price et al., 2005; Seib et al., 2015; Voß et al., 2018). SLPs typically play crucial roles in host-pathogen interactions, including nutrient acquisition, immune evasion, and cell adhesion, indicating that these targets are unlikely to be lost due to selective pressures (Hooda et al., 2017). However, the surface accessibility required to facilitate binding to host ligands inevitably exposes SLPs to the host immune system, often leading to substantial antigenic variation of these proteins (Adamiak et al., 2015; Gandhi et al., 2016). This poses a major challenge for developing SLP-based vaccine compositions with broad coverage.

Despite the inherent variability of these antigens, SLPs have been successfully used in commercial vaccines. Notably, factor H binding protein (fHbp), which helps *Neisseria meningitidis* evade the host complement response by binding the complement inhibitory protein Factor H, is found in two separate commercially licensed protein-based serogroup B meningococcal (MenB) vaccines. To counteract the antigenic variation of this protein, the bivalent MenB vaccine (Trumenba, Pfizer) uses two fHbp variants while the 4CMenB vaccine (Bexsero, GSK) is composed of a single fHbp antigen along with three additional bacterial components to expand coverage (Serruto et al., 2012; Gandhi et al., 2016).

The bipartite bacterial transferrin receptor, which is composed of transferrin binding protein B (TbpB, an anchored SLP) and transferrin binding protein A (TbpA, an integral outer membrane protein), is a highly promising vaccine target with well-established roles in invasive disease and mucosal persistence for a variety of human and animal pathogens from the *Neisseriaceae, Pasteurellaceae*, and *Moraxellaceae* families (Cornelissen et al., 1998; Agarwal et al., 2005; Echenique-Rivera et al., 2011; Pogoutse and Moraes, 2017). This receptor functions for bacterial iron acquisition, where the surface-anchored TbpB extends away from the bacterial surface and binds to iron loaded host transferrin (Tf) (Retzer et al., 1998), followed by removal and internalization of iron across the outer membrane through TbpA (Noto and Cornelissen, 2008). Several studies by our group and others have demonstrated the efficacy of recombinant, purified TbpB or TbpA proteins in mouse immunization and challenge studies against *N. meningitidis* (Nme), the causative agent of invasive meningococcal disease and *Neisseria gonorrhoeae* (Ngo), the causative agent of the sexually transmitted infection gonorrhea. The benefit of targeting these antigens lies in their ability to not only prevent invasive disease, but also potentially extend protection to relevant mucosal surfaces (Lissolo et al., 1995; Price et al., 2005; Fegan et al., 2019). This latter effect is an essential requirement for any gonococcal vaccine in development and would be a significant improvement over the current Group B meningococcal vaccines that have limited impact on nasopharyngeal carriage (Buckwalter et al., 2017; Marshall et al., 2020; McMillan et al., 2021).

Vaccine efforts have largely focussed on TbpB due to its inherent stability, ease of production, and scalability for translation applications. Neisserial TbpBs segregate into distinct lineages, Isotype I and Isotype II, with minimal cross-reactivity or protection being elicited between the two isotypes (Rokbi et al., 2000). The Isotype I lineage is comprised solely of meningococcal TbpBs, while Isotype II can be further divided into four meningococcal and two gonococcal subclusters according to our most recent phylogenetic analysis (Fegan et al., 2025). TbpB consists of two lobes, C-lobe and N-lobe, with the latter binding to the C-lobe of iron loaded (holo) human transferrin (hTf) (Ling et al., 2010). The sequence variability in TbpB across strains mainly resides within the more distal, transferrin-binding interface of the N-lobe, with the β-barrel core of the C-lobe being relatively conserved (Calmettes et al., 2012; Adamiak et al., 2015). We have recently demonstrated that a composition consisting of two rationally selected full-length Ngo TbpBs is sufficient for broad Ngo coverage (Fegan et al., 2025). As the meningococcal TbpBs are more diverse, we postulate that multiple Isotype II variants in combination with an Isotype I TbpB may be necessary for broad meningococcal coverage if a vaccine were to consist of only full-length TbpB antigens. While feasible, multi-antigen vaccines require greater production requirements and result in more expensive end products, thereby limiting accessibility.

In an effort to achieve effective coverage with a minimum number of antigens, our group has engineered a minimized version of an Isotype II Nme TbpB C-lobe, termed the Loopless C-Lobe (LCL). The LCL was generated by deleting the N-lobe sequence and replacing four relatively large, variable loops (26, 30, 21, and 19 amino acids in length) within the C-lobe that were not resolved in the crystal structure of the M982 TbpB (Calmettes et al., 2012) with smaller fragments (2-6 amino acids) from loop regions of *Actinobacillus pleuropneumoniae* TbpB (Moraes et al., 2009). We postulated that the LCL could serve as an antigen with dual modality. First, being an easily scalable, soluble protein, LCL could be utilized as a scaffold to display surface-exposed loops from integral membrane proteins that are otherwise challenging to produce at large scale. Second, by removing the more hyper variable N-lobe and the C-lobe loops, the remaining LCL sequences are highly conserved across the TbpB phylogeny. Therefore, we considered whether immunization with LCL may elicit immune responses to conserved portions of diverse TbpBs, thereby improving coverage.

In support of its utility as a scaffold, we have previously demonstrated that LCL-TbpA hybrids, wherein different surface exposed loops from TbpA were grafted on to LCL, can elicit TbpA-specific antibodies with bactericidal activity and immunization with these immunogens elicited protective immunity in mouse challenge models (Fegan et al., 2019). We have also shown that LCL-based hybrids displaying loops of the zinc-binding protein ZnuD from *Acinetobacter baumannii* can protect against this pathogen in a mouse sepsis model (Qamsari et al., 2020), and the LCL has also been used to display extracellular loops from outer membrane proteins from *Treponema pallidum* (Ferguson et al., 2023). Here we establish the protective capacity of LCL as a stand-alone antigen, demonstrating the extent of anti-LCL antibody coverage of heterologous TbpBs and the efficacy of LCL against heterologous *N. meningitidis* and *N. gonorrhoeae* strains in mouse sepsis and colonization models.

## Results

### Structure and stability of LCL

To create the LCL, we replaced the four native C-lobe loops of M982 TbpB that were not resolved in the full-length crystal structure (PDB 3VE2) (Calmettes et al., 2012) with heterologous linkers. AlphaFold models predict that the deleted loop regions are largely unstructured (**Figure 1A, 1B**). To determine if loop replacements had an impact on the overall structure of the C-lobe core, we solved the crystal structure of LCL (PDB 5KKX) (**Figure 1C**), which demonstrated that the core structure remains intact. Sequence conservation map of the C-lobe, obtained by alignment of 1,485 publicly available TbpB variants, shows that the LCL portion is highly conserved (**Figure 1D**).

**Figure 1.**
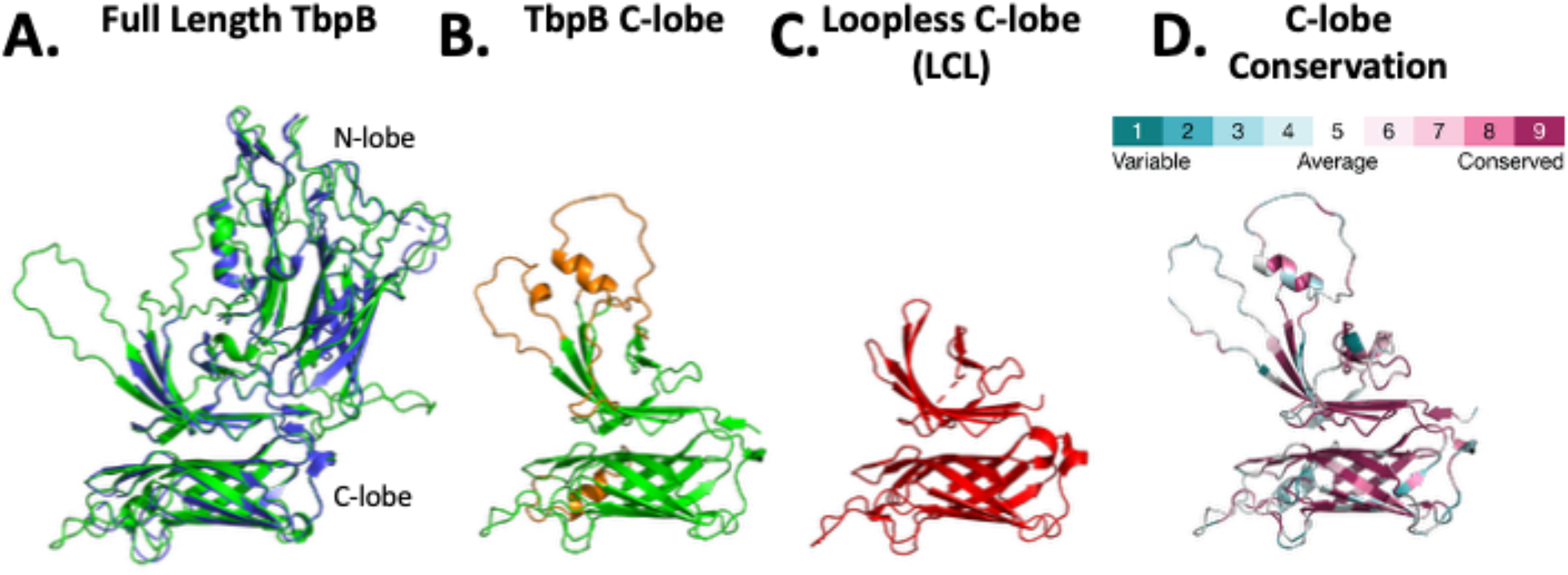
Structural analysis of TbpB and its variants. A: Full-length M982 TbpB structure, solved by x-ray crystallography (PDB 3VE2) (Calmettes et al., 2012), is aligned to the full-length structure predicted by AlphaFold 3. Several large loops are not observed in the crystal structure and are predicted to be disordered. B: The AlphaFold3 predicted structure of M982 TbpB C-lobe, loops that we replaced with shorter heterologous loops in the LCL are shown in orange. C: Structure of the LCL (PDB 5KKX) solved by x-ray crystallography. D: Sequence conservation in the C-lobe calculated using our previous phylogenetic analysis (Fegan et al., 2025) and the ConSurf web server (Yariv et al., 2023).

We next assessed the thermal stability of each recombinant protein (full-length TbpB, C-lobe, and LCL) by measuring the intrinsic fluorescence signal from tryptophan and tyrosine residues using nano differential calorimetry (nanoDSF) (Magnusson et al., 2019), at neutral and acidic pHs to mimic conditions in the extracellular space and the endosome, respectively (**Figure 2A**). Full-length TbpB displayed two transition temperatures (∼50°C and 70°C), which we presume correspond to the unfolding of the N- and C-lobes, as the C-lobe had a single transition (at ∼70°C) that closely matched the second transition of full-length TbpB. Notably, the LCL fluorescence emission ratio at 350nm/330nm increased slightly but did not produce an inflection point, likely due to the β-barrel remaining mostly intact even at elevated temperatures. To further examine the stability of the LCL, proteolytic cleavage by trypsin and chymotrypsin was performed and monitored over time. LCL displayed greater resistance to both enzymes compared to its parental full-length TbpB and C-lobe constructs (**Figure 2B**). Overall, these studies confirm that LCL is a highly stable antigen that closely recapitulates the core tertiary structure of the TbpB C-lobe.

**Figure 2.**
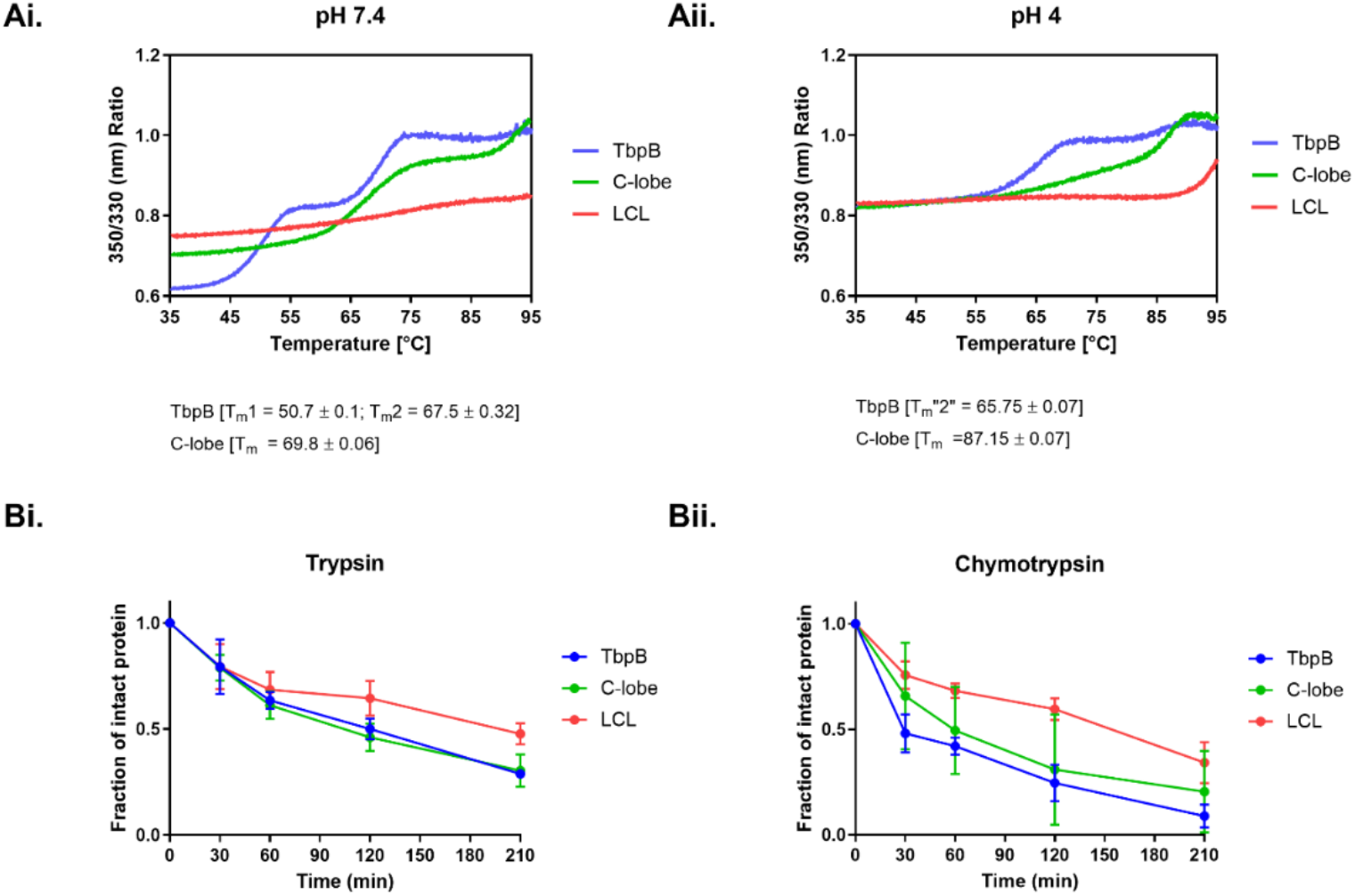
Comparing thermal stability and proteolysis of intact TbpB, C-lobe, and LCL. Ai, Aii: Thermal stability of TbpB, C-lobe and LCL at neutral and acidic pH, respectively, measured using NanoTemper Tycho with the Tm values indicated below. Bi, Bii: Graphs depicting the fraction of intact protein over time after treatment with trypsin and chymotrypsin enzymes, respectively. For trypsin, mean +/- standard deviation shown; n=3 technical replicates per group per time point. For chymotrypsin, mean +/- standard deviation shown; n=4 technical replicates (TbpB and LCL) or n=2 (C-lobe) per time point.

### Immunization with LCL protects against meningococcal sepsis by the homologous Nme M982 strain

To compare the efficacy of LCL to its parental TbpB and full-length C-lobe, C57BL/6 male mice were immunized three times and then challenged with a lethal dose of the homologous *N. meningitidis* M982 strain in a sepsis challenge model (**Figure 3**). As expected, mice that received adjuvant only reached clinical endpoint by 24 hours post infection (0/3 survivors) and were highly bacteremic. All animals immunized with full-length M982 TbpB survived the challenge (4/4 survivors), developed minimal symptoms, and cleared bacteria from the blood by 24 hours post infection. M982 C-lobe immunized group had 50% protection (2/4 survivors), with both survivors clearing bacteria by 48 hours. The loss of efficacy when using individual lobes of TbpB has been a consistent trend in both *N. meningitidis* mouse sepsis studies (**Figure S1A**) and in pigs immunized with full-length TbpB or individual lobes derived from the porcine pathogen *Glaesserella parasuis* (Barasuol et al., 2017).

**Figure 3.**
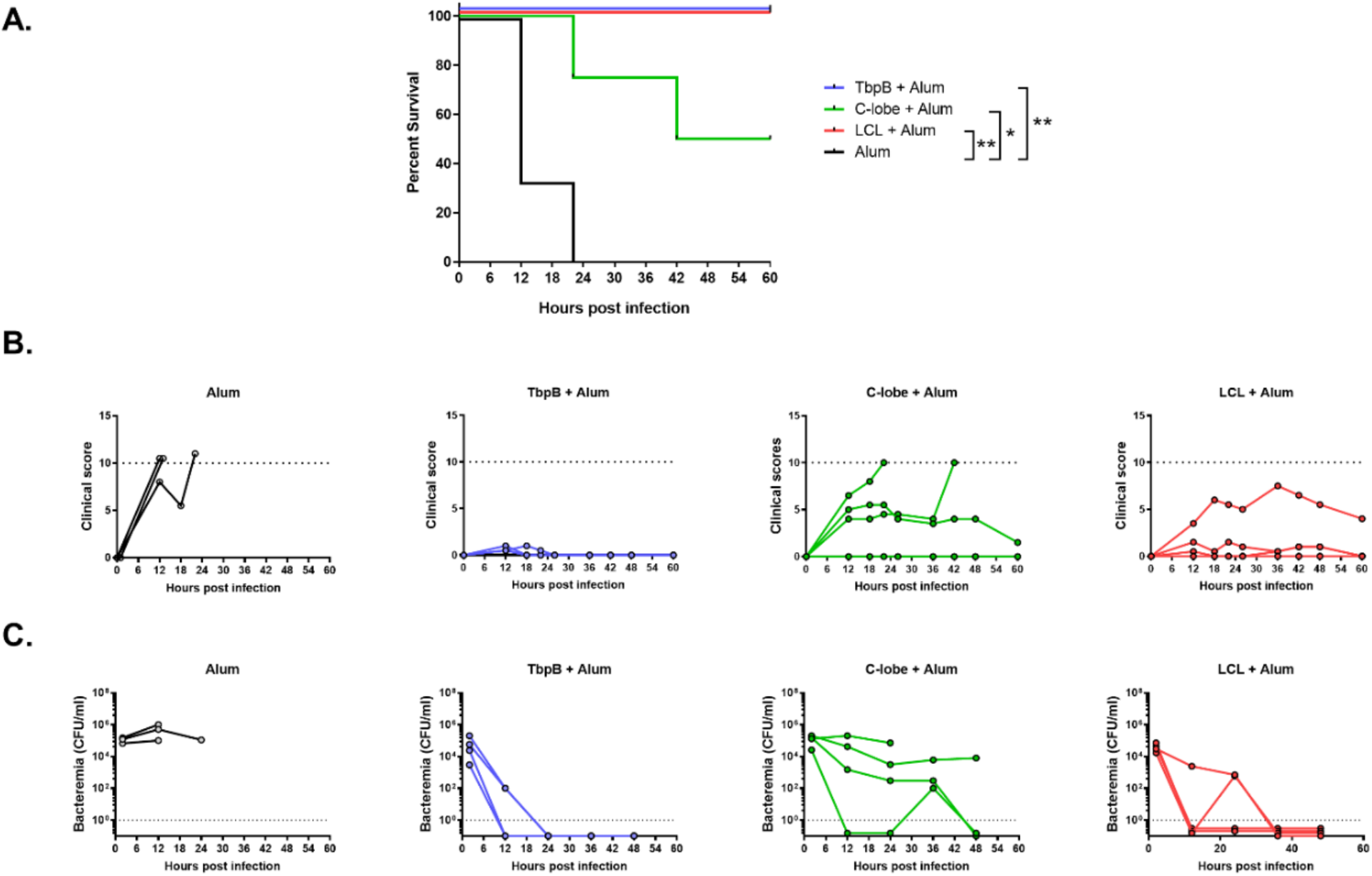
Comparison of TbpB, C-lobe, and LCL to elicit protection against meningococcal sepsis. A: Percentage of survivors after intraperitoneal administration of a lethal dose of the homologous M982 Nme strain. N=3-4 mice per group; p-value calculated using Log-rank (Mantel-Cox) test comparing each immunized group against mice that received alum only; *, p<0.05; **, p<0.01. B: Cumulative clinical score of individual animals within the group indicated above each graph. Dotted line at 10 depicts the clinical score cutoff for humane endpoint. C: Bacterial recovery from tail vein bleeds from the time points indicated. Dotted line at 1 depicts the detection limit. Statistical analysis performed using GraphPad Prism 10.2.0.

Strikingly, mice that received LCL were fully protected from this challenge (4/4 survivors). Clinical scores showed 3 out of 4 mice were protected from clinical symptoms, with the one symptomatic mouse recovering by the end of the study. 2 out of 4 mice cleared bacteria within 12 hours and the remaining animals cleared by 36 hours post infection. These results demonstrate that LCL is a superior antigen to the C-lobe and can yield comparable protection to full-length TbpB against the homologous strain during invasive challenge.

### LCL protects against nasopharyngeal colonization by the homologous Nme M982 strain

Next, we utilized human carcinoembryonic antigen-related cell adhesion molecule 1 (hCEACAM1)-expressing transgenic mice which are permissive to Nme colonization of the nasopharynx after direct instillation of bacteria at the nares (Johswich et al., 2013). Using this model, we have previously reported that the 4CMenB vaccine can reduce nasopharyngeal carriage of only a subset of single-antigen matched meningococcal strains, suggesting that not all antigens are able to provide adequate mucosal protection after parenteral vaccination (Buckwalter et al., 2017). In direct support of this premise, comparison of fHbp to full-length TbpB revealed that while both fully protect against sepsis and elicit significant bactericidal titres (**Figure S1A, C**), only the latter reduces nasal colonization rates (from 64% for adjuvant and 58% for fHbp groups to 18% for the TbpB-immunized group, **Figure S1B**). Therefore, we proceeded to evaluate if LCL can retain this efficacy of full-length TbpB.

Full-length TbpB, C-lobe, and LCL-Immunized hCEACAM1 FvB mice were intranasally challenged with the homologous Nme M982 strain and the bacteria recovered from the nasopharyngeal tissue was enumerated three days post infection. The percentage of culture positive animals in the adjuvant control group was 47% (7/15 mice), while full-length TbpB, C-lobe, and LCL-immunized groups had a drop in the colonization rates to 18.2% (2/11), 28.6% (2/7), and 11.2% (1/9) respectively (**Figure 4A**). Since M982 is not a robust colonizer of the murine nasopharynx and to ensure reproducibility, we performed three additional independent studies comparing only LCL to adjuvant in hCEACAM1 transgenics from different genetic backgrounds and varying routes of immunization. In each of these studies, the colonization rate in LCL-immunized animals were consistently lower than adjuvant controls: LCL+Alum – 20% (1/5), 33% (2/6), 25% (2/8) versus Alum – 80% (4/5), 50% (4/8), 62.5% (5/8) respectively, although the reduction in bacterial burden did not reach statistical significance in these individual experiments (**Figure 4B, C, D**). These results demonstrate that parenteral immunization with LCL provides consistent reduction in nasal colonization rates, with its performance being comparable to full-length TbpB against the homologous challenge strain, Nme M982.

**Figure 4.**
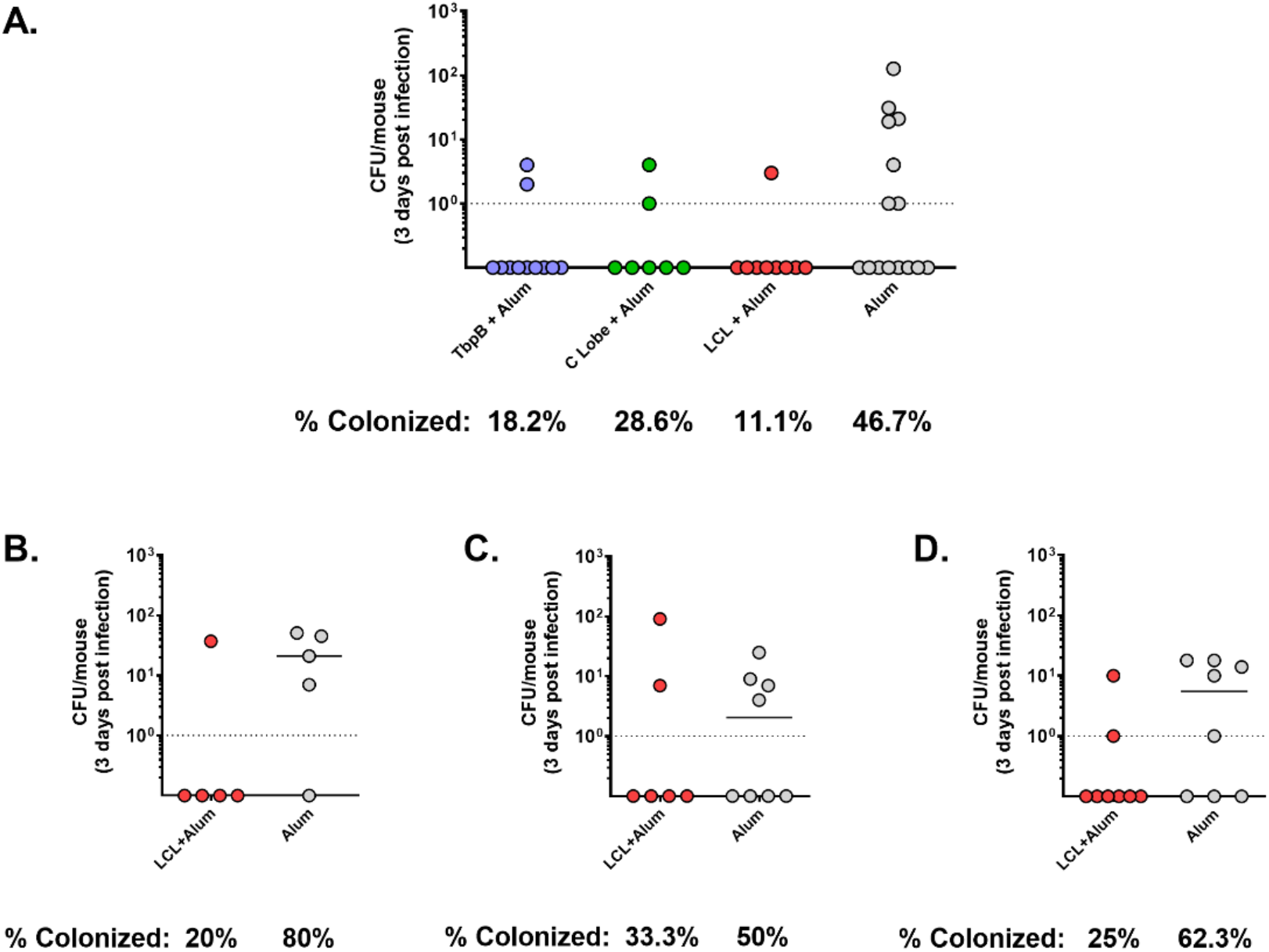
LCL-mediated protection against nasal colonization by the homologous Nme M982 strain. Meningococcal recovery from the nasopharynx of immunized hCEACAM1 transgenic mice 3 days post nasal administration of the homologous M982 Nme strain, with the percentage of culture positive animals indicated below. A: Comparing protection mediated by TbpB, C-lobe, and LCL intraperitoneal immunization in hCEACAM1+/- (mCEACAM1+/+) FvB mice. N=7-15 for each group. B-D: Comparing LCL to adjuvant control in 3 additional independent experiments with the following conditions: B: Intraperitoneally immunized hCEACAM1+/- (mCEACAM1+/+) FvB mice; C: subcutaneously immunized hCEACAM1+/- (mCEACAM1-/-) C57BL/6 mice; D: intraperitoneally immunized hCEACAM1+/- (mCEACAM1-/-) C57BL/6 mice. N=5-8 for each group. Each circle represents Nme recovered from one mouse, line at median. One-way ANOVA with Dunnett’s multiple comparison of the bacterial burden in each vaccinated group to the control (A) and non-parametric Mann-Whitney test (B-D) performed using GraphPad Prism 10.2.0 did not yield significant p-values.

### LCL elicits robust levels of functional IgG

Protection against invasive meningococcal infection is antibody-dependent (Borrow et al., 2005). Considering that LCL and full-length TbpB, but not C-lobe, fully protected in the sepsis model, we measured anti-TbpB serum IgG levels among these groups. Pre-challenge serum from the sepsis study in Figure 2 and terminal serum from the colonization study in Figure 3A were assayed using protein-based and heat inactivated whole bacterial ELISAs (**Figure 5A-C, D** respectively). As expected, immunization with full-length TbpB resulted in the highest antibody titre against the full protein and whole bacteria, with recognition of both C- and N-lobes, whereas immunization with the C-lobe resulted in significantly lower anti-TbpB antibodies and no N-lobe reactivity. Compared to C-lobe, immunization with LCL yielded significantly higher IgG titres in protein-based ELISAs (**Figure 5A, B**), implying that the latter is more immunogenic. In contrast, the difference in titres against whole bacteria between LCL and C-lobe sera was subtle (**Figure 5D**), presumably due to proximity of other membrane proteins and/or the polysaccharide capsule limiting antibody access, or artifacts arising during whole bacterial plate preparation masking some epitopes.

**Figure 5.**
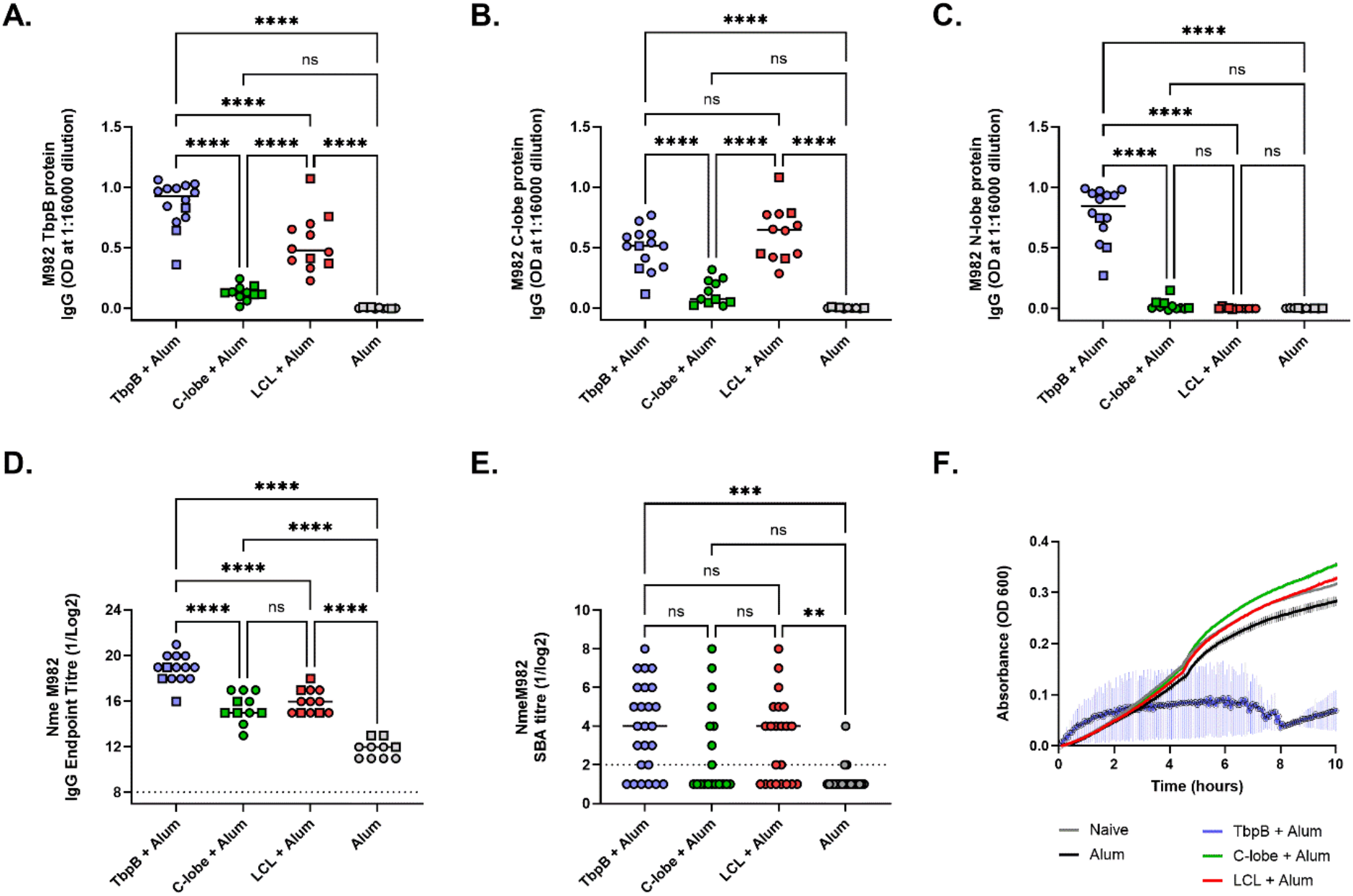
Comparison of anti-TbpB antibodies elicited by TbpB, C-Lobe, and LCL antigens. A-C: Protein ELISA measuring reactivity of TbpB, C-lobe, and LCL-immune sera against full-length TbpB, TbpB C-lobe, and TbpB N-lobe, respectively. D: IgG titre elicited by the different immunogens towards heat inactivated Nme M982. A-D: N=10-15 per group; squares represent pre-challenge sera from C57BL/6 animals from the sepsis study presented in Figure 2; circles represent terminal serum from hCEACAM1 FvB mice from the colonization study presented in Figure 3A, each symbol represents serum from one animal tested in duplicate. E: Serum bactericidal activity (SBA) titre against iron-starved Nme M982. N=19-25 per group, including terminal hCEACAM1 FvB serum from Figure 4A, plus additional terminal serum from immunized hCEACAM1 FvB from pilot studies. SBA titre (>50% killing) reached for 19/25 (76%) TbpB, 8/19 (42%) C-lobe, 14/22 (64%) LCL and 3/21 (14%) Alum samples. Dotted lines represent lowest dilution tested. Line at median for each group. One-way ANOVA with Tukey’s post-hoc test comparing each group to every other group performed using GraphPad Prism 10.2.0. For D, E: Statistics performed on Log2 transformed data. ns, not significant; **, p<0.01; ***, p<0.001; ****, p<0.0001. F: Meningococcal growth inhibition assay. Iron-starved M982 strain was grown in the presence of hTf and pooled heat-inactivated serum from TbpB, C-lobe and LCL immunized animals from Figure 4A. Change in absorbance (OD600) relative to the initial time point is graphed; error bars depict standard deviation of two technical replicates.

Serum bactericidal activity has long served as a reliable correlate of protection against invasive meningococcal disease and is considered the gold standard in the field (Borrow et al., 2005). Bactericidal titre also correlated with protection in our comparative study of fHbp and TbpB antigens (**Figure S1C**, only the groups that were protected in the sepsis study in **Figure S1A** had significant bactericidal titres). Therefore, to elucidate antibody-dependent mechanisms contributing to LCL-mediated protection, we next performed serum bactericidal assays. LCL serum had significant bactericidal activity as did full-length TbpB serum, while C-lobe serum lacked activity (**Figure 5E**), which reflected the protection outcome in the sepsis challenge. We additionally compared the ability of these sera to directly block human transferrin dependent bacterial growth. With full-length TbpB antigens, we consider this additional mechanism of nutritional deprivation by antibodies competing for the transferrin binding site on TbpB important for mediating protection (Frandoloso et al., 2015). Although LCL antibodies do not bind the N-lobe of TbpB where the transferrin-binding interface lies, we wondered whether antibodies binding to the C-lobe could adect transferrin utilization indirectly by hindering either TbpB-hTf and/or TbpB-TbpA interactions. To test this, we compared the growth of M982 Nme in the presence of serum and human transferrin. Only TbpB serum, but not LCL or C-lobe serum was able to inhibit transferrin dependent bacterial growth (**Figure 5F**). Taken together, these results suggest that LCL antibodies facilitate bacterial clearance through complement-mediated bacterial lysis, but not nutrient starvation.

### Anti-LCL antibodies are highly cross-reactive

Since the LCL was engineered to remove most of the hypervariable sequences within TbpB, we next examined whether this strategy results in broadly cross-reactive antibodies being raised against conserved regions of TbpB. Phylogenetic analysis of all publicly accessible TbpB sequences results in the sequences dividing into several clusters, with the meningococcal Isotype I TbpBs forming a single cluster (Isotype I) and Isotype II TbpBs separating into four Nme (Nme 1, 2, 3, 4) and two Ngo clusters (Ngo 1, 2) (**Figure 6A**). To evaluate cross-reactivity of immune sera, we first selected representative TbpBs from different clusters (N=3 from Nme 1; N=6 from Nme 2, including the homologous M982 TbpB; N=3 from Nme 3; N=2 from Ngo 1; N=2 from Ngo 2, N=1 from Isotype I) for high throughput protein ELISAs (Fegan et al., 2021). Signal obtained against each TbpB is depicted as a heat map to showcase individual responses and as bar graphs for overall group trends (**Figure 6B, C**, respectively). Signal from HRP-labelled hTf (hTf-HRP) were used to confirm uniform coating of the ELISA plate with properly folded analyte (**Figure 6C, inset**).

**Figure 6.**
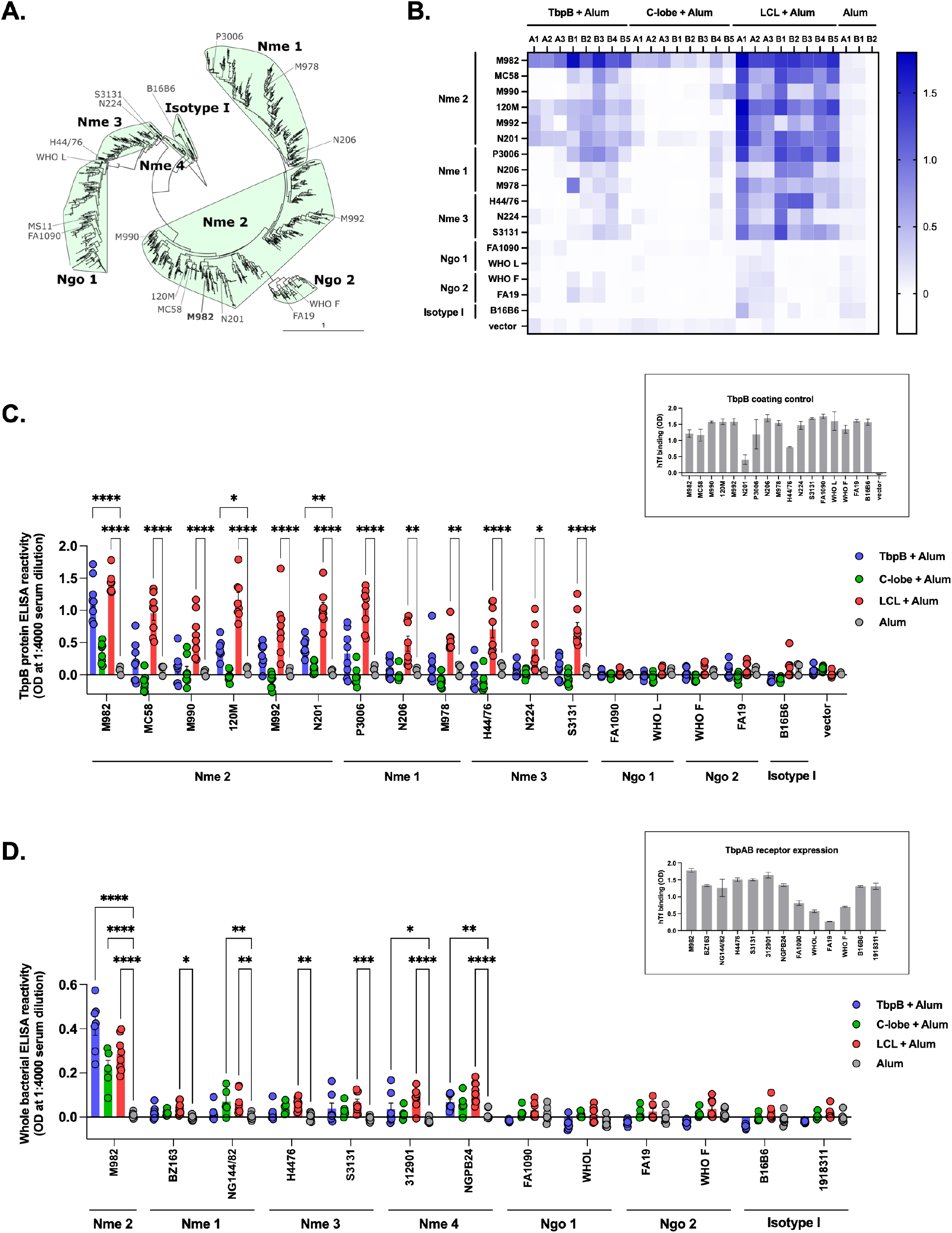
Antibody cross-reactivity against TbpB proteins. A: Phylogenetic tree depicting Neisserial TbpB diversity. The location of M982 TbpB is highlighted, along with 16 additional TbpBs spanning various clusters used for cross-reactivity analysis. B: Heat map depicting IgG cross-reactivity against a broad panel of TbpB protein variants by ELISA. Optical density (OD) readings denoted by the colour gradient. Background noise from no-protein control subtracted. Each row is a different TbpB protein capture, with the phylogenetic cluster indicated to the left. Each column is a different mouse sample, with the immunizing antigens indicated above. Numbered labels starting with A are pre-challenge serum from the immunized C67BL/6 sepsis cohort; B are terminal serum from the immunized hCEACAM1 FvB colonization cohort. C: Grouped bar graph summarizing cross-reactive antibody data from B. Bars represent mean, error bars represent standard error, each circle represents serum from a single mouse (N=8 for immune sera, N=3 for Alum). hTf-HRP signal quantifying properly folded TbpB capture is depicted in the inset, mean ± standard deviation. D: Serum reactivity against representative Nme and Ngo strains grown under iron limitation using inactivated whole bacterial ELISA. TbpB cluster indicated below the x-axis. Background noise from no serum control subtracted. Bars represent mean, error bars represent standard error, each circle represents serum from a single mouse, N=5-8/group. Inset depicts hTf-binding as an indicator of TbpAB receptor expression on bacteria used for coating ELISA plates, mean ± standard deviation. For C and D, two-way ANOVA with Dunnett’s post-hoc test comparing each group to Alum control performed using GraphPad Prism 10.2.0. Only p-values <0.05 shown. *, p<0.05; **, p<0.01; ***, p<0.001, ****, p<0.0001.

Despite high reactivity against the homologous TbpB, serum from full-length TbpB-immunized animals reacted weakly against heterologous TbpBs. Reactivity varied based on mouse background, with C57BL/6 serum only recognizing a subset of Nme 2 TbpBs and hCEACAM1 FvB serum recognizing additional Nme 1 and Nme 3 TbpBs (**Figure 6B**). Overall, significant cross-reactivity with anti-TbpB IgG was only observed against two Nme 2 TbpBs: 120M and N201 (**Figure 6C**). C-lobe sera reacted primarily to the homologous M982 TbpB, with only a single hCEACAM1 FvB sample producing a broader reactivity pattern. Remarkably, serum from LCL-immunized animals recognized all Nme Isotype II TbpBs in our panel. This LCL-elicited response was significant and consistent regardless of mouse background.

Next, we examined cross-reactivity against a panel of Nme and Ngo strains from all heterologous TbpB clusters (N=2 strains from Nme 1, 3, 4, Ngo 1, 2, Isotype I; N=5-8 for each serum group) (**Figure 6D**; inset depicts hTf-binding as an indicator of transferrin receptor surface expression, however this control cannot discern between TbpA and TbpB binding). Consistent with the pattern observed with protein-based ELISAs, anti-LCL antibodies significantly cross-reacted against all Nme Isotype II strains, increasing strain coverage compared to full-length TbpB antigen which only had significant cross-reactivity signal against strains from the Nme 4 cluster. We further performed western blots on the same strain panel to confirm that LCL serum is specifically recognizing TbpBs (**Figure S2**).

While characterization of these cross-reactive antibodies is beyond the scope of this study, the consistent recognition of Isotype II Nme TbpBs, but not Ngo TbpBs, leads to the consideration of whether there are regions within the LCL that are conserved in Nme TbpBs, and distinct in Ngo TbpBs, that could provide clues regarding the location of potential cross-reactive epitopes.

### LCL immunization improves clinical outcome during invasive infection by heterologous Isotype II Nme strains

We have previously observed that passive transfer of anti-LCL immune serum is protective against invasive challenge and colonization by the homologous strain (**Figure S3**), providing direct evidence of antibody-mediated protection in these infection models. This, combined with the broad strain recognition by anti-LCL antibodies (**Figure 5**) prompted us to conduct challenge studies with strains from each heterologous Nme Isotype II (Nme 1, Nme 3, Nme 4) and Isotype I clusters to assess the breadth of cross protection. LCL-immunized C57BL/6 mice were intraperitoneally infected with 10^6^-10^7^ colony forming unit (CFU) of the selected Nme strains and then monitored for the progression of clinical symptoms (**Figure 7**), with mice reaching a clinical score of 10 being considered at clinical endpoint and humanely euthanized. Consistent with antibody coverage, clinical scores in mice immunized with LCL with all three Isotype II Nme strains (NG144/82 (Nme 1), H44/76 (Nme 3), NGPB24 (Nme 4), having TbpBs sharing 82%, 68% and 69% sequence identity with full-length M982 TbpB, respectively) were lower than the clinical score for mice that received only the adjuvant control. There was no difference in clinical outcome when mice were challenged with the Isotype I Nme strain (B16B6, 37% sequence identity with M982 TbpB), consistent with the lack of antibody cross-reactivity seen.

**Figure 7.**
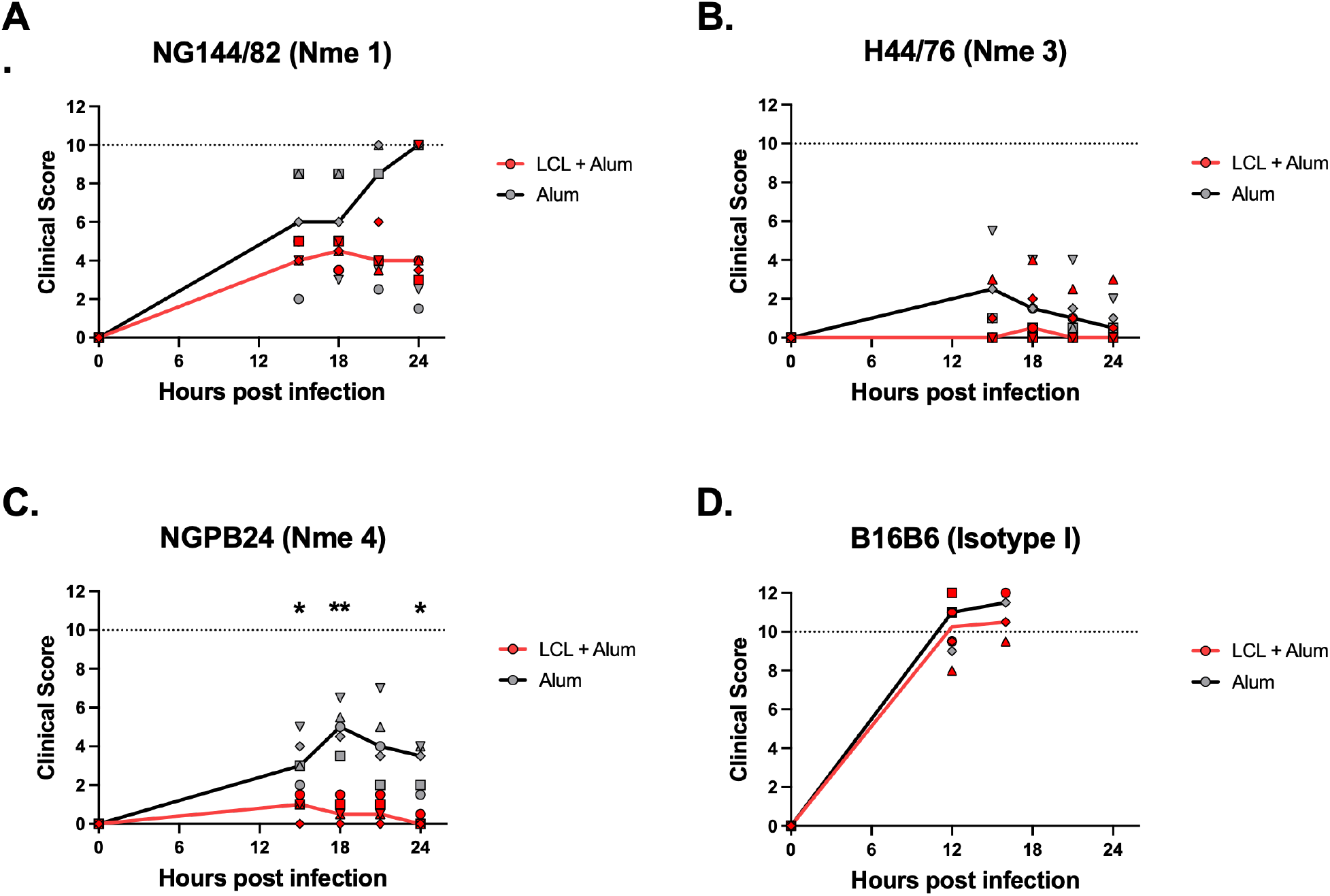
Cross-protection against invasive disease by heterologous Nme strains. Graphs showing total clinical score after intraperitoneal injection of the indicated *N. meningitidis* strain into LCL-immunized (red) or control (grey) C57BL/6 male mice. N=5 mice per group; with each mouse depicted by a different shape. Thick solid line connects median clinical score at each time point. Dotted line at 10 shows the clinical score cutoff at which animals were humanely euthanized. Two-way ANOVA with Bonferroni’s multiple comparison test comparing vaccinated versus control groups at each timepoint was performed using GraphPad Prism 10.2.0. Only p-values <0.05 indicated. *, p<0.05; **, p<0.01.

### LCL protects against nasal colonization by heterologous Isotype II strain

Finally, we assessed LCL-mediated cross-protection against mucosal colonization. Vaccinated hCEACAM1 transgenic mice were intranasally challenged with an Isotype II strain S3131 (**Figure 8A**) that expresses a TbpB from Nme cluster 3 with 69% amino acid sequence identity to M982 TbpB. Bacterial burden was quantified in nasopharyngeal tissues 3 days post infection, revealing the proportion of culture positive animals as the primary readout. Compared to adjuvant alone (55% or 6/11 colonized), the homologous S3131 TbpB antigen (included as a positive control) was sterilizing (0% or 0/8) against this strain. LCL immunization reduced the colonization rate compared to adjuvant alone (33% or 3/9), demonstrating its ability to provide partial mucosal cross-protection.

Since mechanisms underlying mucosal protection are not yet well-defined, despite the absence of LCL mediated antibody cross-reactivity to gonococcal TbpBs and meningococcal Isotype I TbpBs, we proceeded to evaluate nasopharyngeal protection by a TbpB Isotype I Nme strain (90/18311; 37% sequence identity to M982 TbpB) in transgenic hCEACAM1 mice and lower genital tract protection against an Ngo 1 (FA1090; ∼60% sequence identity) and an Ngo 2 strain (FA19; ∼68% sequence identity) in wild type female C57BL/6 mice. In these studies we found no difference in mice immunized with LCL compared to those that received the adjuvant only control (**Figure 8B, Figure S4**).

**Figure 8.**
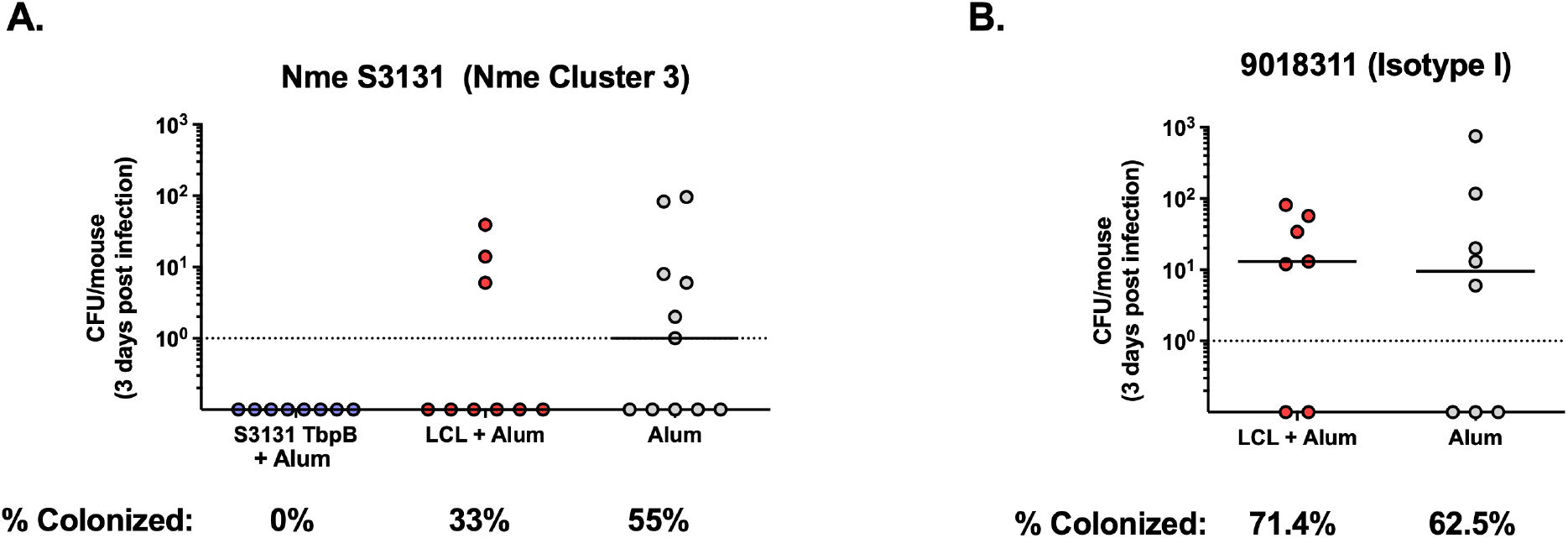
Cross-protection against colonization by heterologous Nme strains. Meningococcal burden in the nasopharynx three days post nasal administration of A: Nme 3 Isotype II strain S3131 and B: Isotype I strain 90/18311. The percentage of culture positive animals indicated below. N=7-11 hCEACAM1+/- (mCEACAM1+/+) FvB mice in each group. Each circle represents CFU from one animal, line at median. One-way ANOVA with Dunnett’s multiple comparison of the bacterial burden in each vaccinated group to the control (A) and non-parametric Mann-Whitney test (B) performed using GraphPad Prism 10.2.0 did not yield significant p-values.

Together, our cross-protection studies indicate that in addition to protecting against invasive infection, LCL may reduce colonization rates by Isotype II Nme strains.

## Discussion

The engineered LCL presented here is comprised of a minimized structure with the conserved regions of the candidate vaccine antigen, TbpB, towards the development of a broadly protective vaccine against the pathogenic *Neisseria* species. While originally developed as a soluble scaffold to display loops from integral membrane proteins that are otherwise not viable for commercial production, in this study we show that LCL alone exhibits remarkable stability and functionality as an independent immunogen, suggesting its potential as a versatile tool for vaccine applications.

The fact that LCL was able to protect against homologous challenge to a similar extent as full-length TbpB was unexpected. Our earlier studies comparing full-length TbpB to C- or N-lobes (Figure S1) suggested that conformational antibodies near the junction of the two lobes may be critical since protection was lost when individually produced lobes were combined. Furthermore, studies in pigs using engineered full-length TbpB antigens indicated that N-lobe specific antibodies that block host transferrin from binding TbpB on the surface of *G. parasuis* during infection are critical for protection.

This is likely due to iron starvation caused by antibodies that target the hTf binding site that provide an added protective mechanism (Frandoloso et al., 2015). Therefore, the ability of LCL to retain protective function despite its significant truncation and inability of elicit antibodies that block transferrin mediated iron uptake provides new insights into protection determinants.

Protein ELISAs clearly demonstrated that anti-LCL antibodies bind the C-lobe specifically, and do not cross-react with the N-lobe, implying that effective targeting of the C-lobe alone can be sudicient to elicit protection. That raises further questions as to why immunization with LCL, but not C-lobe, was effective. One explanation could be that the increased fold stability and proteolytic resistance of LCL allows this antigen to retain its tertiary structure after immunization, eliciting robust conformational antibodies, as reported in case of other stabilized immunogens (Delamarre et al., 2006; Joyce et al., 2016). Differences in performance may also lie in differences in what antibody repertoire is elicited between C-lobe and LCL. The sequence variability of the C-lobe loop regions suggest that the loops are immunogenic and have been subjected to selective pressures over time. Thus, it is reasonable to conclude that anti-C-lobe antibodies may largely be against the loop regions and thus distinct from the LCL repertoire. While beyond the scope of this study, comparison of T cell responses would also be required for fully appreciating the immunological differences.

Our initial hypothesis that immunization with LCL could result in broader TbpB coverage compared to the parental full-length M982 TbpB was also confirmed in this study. Anti-LCL antibody specificity for diverse Nme Isotype II TbpBs, but not Ngo TbpBs that cluster within the overall Isotype II diversity, was somewhat unexpected considering that the Ngo 2 subcluster, in particular, branches off the larger Nme 2 subcluster to which M982 TbpB belongs. However, this specificity provides an opportunity for future work to identify residues that align within LCL that are unique to Nme Isotype II TbpBs and map potential cross-reactive and cross-protective B cell epitopes. This would be crucial in guiding the future design of LCL-based hybrid antigens to avoid disrupting its inherent protective capabilities.

A distinguishing feature of full-length TbpB and its derivative antigens is its ability to protect against mucosal colonization (Fegan et al., 2025; Islam et al., 2025). Interestedly, this is not a common feature of all SLPs since immunization with M982 fHbp against the homologous strain in this study or with 4CMenB vaccine against the fHbp-variant matched strain in a previous study (Buckwalter et al., 2017) did not reduce meningococcal colonization rates in our hands in transgenic hCEACAM1 mice, and the 4CMenB vaccine does not protect against Nme colonization in humans (McMillan et al., 2022). This discrepancy is likely due to an interplay of both host responses to the different vaccine antigens and bacterial responses to different environments regulating expression of these targets. The mucosal protection conferred by TbpB-specific antibodies may, therefore, result from the combination of their ability to starve the bacteria of iron and clear the bacteria by opsonization-dependent processes. As the work presented here was performed in mice lacking human transferrin, the contribution of iron restriction on meningococcal clearance from mucosal surfaces remains to be elucidated.

Overall, SLPs have been recognized as enticing candidate vaccine antigens for a wide variety of gram-negative pathogens. SLPs are consistently comprised of a β-barrel and a handle domain, a pattern duplicated in each lobe of TbpB. The efficacy shown here of a minimized version of a single lobe of TbpB offers a framework for developing other SLPs as more broadly cross-protective antigens by removing hypervariable loops while focusing on maintaining antigen stability. As more SLPs are identified in pathogens of interest, this strategy of antigen engineering may allow for accelerated vaccine development going forward.

Together, we have demonstrated that engineering a meningococcal TbpB to a minimized, stable LCL provides a superior immunogen that elicits a broadly cross-protective immune response and presents a novel strategy towards developing broad-spectrum protein-based vaccines.

## Materials and Methods

### Protein production

Full-length meningococcal M982 TbpB, C-lobe, and LCL were cloned into a custom T7 protein expression vector, downstream and in fusion with an N-terminal polyhistidine tagged maltose binding protein (MBP) followed by a TEV cleavage sequence as previously described (Fegan et al., 2019). Protein expression vectors were heat-shocked into chemical competent *E*.*coli* T7express cells (New England Biolabs). The transformants were allowed to recover in LB broth for 1 hour, shaking at 175 RPM and 37°C, before selection by additional LB supplemented with ampicillin (100 μg/mL) and grown for a further 3 hours. The starter culture was then used to inoculate 6 L of autoinduction ZY media (Studier, 2014) with 50 μg/mL of ampicillin. The culture flasks were shaken overnight at 37°C and 175 RPM for 16 hours and a further 24 hours at 20°C. Cells were harvested by centrifugation at 5,000 x g for 30 minutes at 4°C. Cells were lysed by homogenization (EmulsiFlex-C3, Avestin) supplemented with protease inhibitor (c0mplete™ Mini, Roche), lysozyme, and DNase I. The resulting lysate was clarified by centrifugation at 16,000 x g for 90 minutes at 4°C, followed by syringe filtration (0.22 μm). Clarified lysate was then recirculated by a peristaltic pump and affinity captured on HisTrap EXCEL columns (Cytiva) overnight, which was then mounted on an ÄKTA Purifier 100 (GE Healthcare). The column was washed and protein eluted by imidazole, and the eluted fraction was dialyzed overnight into a low imidazole buffer to facilitate cleavage by the addition of in-house purified TEV protease to separate the N-terminal MBP fusion partner. The cleaved protein was then exchanged into a low salt buffer and loaded onto an anion exchange HiTrapQ column (Cytiva), and the fractions containing TbpB, C-lobe or the LCL were concentrated and passed through a size exclusion column (HiPrep 26/60 Sephracryl S-200 HR, Cytiva) to further remove any contaminants. A final polishing step with a strong anion exchange MonoQ 5/50 GL column (Cytiva) was used to remove lipopolysacchrides, the final samples were concentrated, and aliquots were flash frozen by liquid nitrogen. Sample purity was assessed by 12% SDS-PAGE and protein concentration was determined by NanoDrop (ThermoFisher).

### Crystallization of LCL

Purified LCL was initially screened with an automated Gryphon robot (Art Robbins) against Hampton and JCSG+ (Qiagen) commercial screening suites using sitting drop vapour diffusion. Initial hits were optimized with a 1:1 (protein: precipitant) ratio in a precipitant solution composed of 0.1 M Bicine pH 9.0, 20% PEG 6000 that yielded crystals that diffracted to 1.95 Å in space group P 4 21 2.

### Data collection and structure determination

Crystallographic data was collected on crystals at 80 K at the Advanced Photon Source (NECAT-24-ID-E beamline). The diffraction dataset was collected at a wavelength of 0.9795 Å and processed using DENZO and SCALEPACK from the HKL-2000 suite (Otwinowski and Minor, 1997). The first structural model for the LCL was obtained by molecular replacement using Phenix PHASER (Bunkóczi et al., 2013) with a starting model based on a C-lobe truncation from the full-length M982 TbpB crystal structure (PDB 3VE2). The final model was generated following several rounds of model building and refinement using Coot (Emsley and Cowtan, 2004) and Phenix REFINE (Adams et al., 2010). Data refinement and statistics are summarized in Table S1.

### Accession codes

Structure factors and atomic coordinates for M982 TbpB LCL have been deposited to the PDB under accession code 5KKX.

### Nano differential scanning fluorimetry

Measurements were carried out on a Tycho NT.6 (NanoTemper). Purified protein was diluted to a final concentration of 0.2 mg/mL in either high pH (50 mM sodium phosphate, 150 mM NaCl, pH 7.5) or low pH (50 mM sodium citrate, 150 mM NaCl, pH 4.0) buffers. Samples were loaded into a glass capillary through capillary action and then placed over the instrument mirror. Measurements were conducted between 35°C and 95°C at a rate of 30°C/min.

### Protease stability assay

Antigen susceptibility to proteases was assessed using trypsin (Promega) and chymotrypsin (Bioshop) at protease:antigen ratios of 1:1000 and 1:100 respectively. A 60 μL solution of 0.2 mg/mL purified protein was co-incubated with either protease at 37°C for 0.5 h to 4 h, taking samples at indicated time points. For each time point, 10 μL of reaction was quenched by the addition of 5mM PMSF and mixed with 10 μL of SDS loading buffer and boiled for 5 min. Samples were run on a 15% SDS-PAGE gel and subsequently assayed with mass spectrometry to identify proteolytic fragments.

### Mouse immunizations

Purified protein was first diluted in phosphate buffered saline (PBS) and then mixed with 2% Alhydrogel to a final concentration of 25 μg protein and 100 μg Alum per 100 μL dose and allowed to adsorb for at least 20 mins with gentle agitation. For some studies, Emulsigen D (MVP adjuvants) ready-to-use emulsion was added to diluted protein to a final concentration of 20% (v/v) per 100 μL dose. Vaccines were administered on Days 0, 21, and 42 via intraperitoneal injection, unless otherwise specified in the figure legend. For passive immunizations, terminal serum from a New Zealand White rabbit immunized 3 times subcutaneously with 50 μg of LCL with 20% (v/v) Emulsigen D was obtained from the Schryvers lab (performed under protocol AC11-0033, approved by the University of Calgary Animal Care Committee). Serum was heat inactivated at 56°C for 30 minutes and 250 μL was injected intraperitoneally 6 hours prior to infection.

### Invasive meningococcal mouse infections

4-6 week old male C57BL/6 mice were purchased from Charles River and vaccinated after a minimum of seven days of acclimatization to the facility. Infections were performed approximately 2 weeks after the final vaccine dose. Overnight lawn of the indicated Nme strain was grown on GC-Isovitalex (BD cat. 11875) or GC-Kellogg’s and sub-cultured in RPMI (Wisent, cat no. 350-000) for approximately 3-4 h at 37°C at 120 rpm. Bacteria were diluted in PBS containing calcium and magnesium (PBS++) and a 250 μL suspension containing approximately 10^6^-10^7^ CFU was intraperitoneally injected into isoflurane anaesthetized mice. 8 mg of holo human transferrin (Sigma, cat no. T4132) resuspended in 250 μL of PBS was administered at the same time as infection by a separate intraperitoneal injection for all studies, except Figure S1A where 2 mg of iron dextran in the same volume was used. Tail vein blood was sampled to monitor bacterial load in some studies and enumerated on GC Isovitalex with VCNT inhibitor (vancomycin, colistin, nystatin, and trimethoprim for selection of Neisseria species; BD BBL™ VCNT Inhibitor (Thayer and Martin), Fisher Scientific, cat. B12408). Clinical symptoms were scored at regular intervals and a cumulative score of 10 or a weight loss of 20% was used as the clinical cutoff for humane euthanasia by cervical dislocation. Studies were performed under the animal use protocol 20011319, approved by the Animal Care Committee at the University of Toronto. For all mouse studies, mice were kept under specific pathogen free conditions and given access to food and water *ad libitum*.

### Meningococcal nasopharyngeal colonization

Approximately 6 week old transgenic hCEACAM1+/- (mCEACAM1+/+) FvB and hCEACAM1+/- (mCEACAM1-/-) C57BL/6 mice bred in-house were immunized and then challenged intranasally approximately 2 weeks after the final dose. Briefly, overnight lawn of mouse passaged Nme grown on GC-Isovitalex (BD cat. 11875) at 37°C with 5% CO2 was suspended in PBS++ and approximately 10^7^ CFU in 10 μL was inoculated into the nostrils of isoflurane anaesthetized mice. Mice were humanely euthanized by CO2 overdose 3 days post infection. Terminal serum was collected by cardiac puncture and stored at -20°C until analysis. Retrograde nasal lavages were collected by injecting 250 μL PBS++ through the trachea and out the nostrils and nasal turbinate was swabbed. Lavage and swab samples were grown on GC-Isovitalex-VCNT plates for 48 h at 37°C with 5% CO2 for selection and enumeration of *N. meningitidis*. Meningococcal colonization studies were performed under the animal use protocol 20011319, approved by the Animal Care Committee at the University of Toronto.

### Gonococcal lower genital tract colonization

Approximately 4-6 week old female C57BL/6 mice were purchased from Charles River and vaccinated after a minimum of seven days of acclimatization to the facility. Approximately 2 weeks after the final dose, estrous cycle was monitored, hormones and antibiotics were administered and the mice were infected with Ngo as described previously (Islam et al., 2025). To prepare the inoculum, overnight lawns of Ngo FA1090 and FA19 grown on GC-Isovitalex were suspended in RPMI and ∼5 μL containing approximately 10^7^ CFU was vaginally instilled. Bacteria were enumerated from vaginal lavage samples as previously described (Islam et al., 2025) to determine colonization duration. Upon study completion, mice were humanely euthanized by CO2 overdose followed by cervical dislocation. Gonococcal studies were performed under the animal use protocol 20011775, approved by the Animal Care Committee at the University of Toronto.

### ELISAs

For protein ELISAs, 384 well plates (VWR, cat. CA62409-064) were coated with TbpB as previously described (Fegan et al., 2025). For whole bacterial ELISAs, strains were grown overnight on GC-Isovitalex containing 10 μM deferoxamine mesylate salt (Sigma, cat. D9533), restreaked onto fresh GC-Isovitalex-10 μM deferoxamine plates for another overnight growth at 37°C with 5% CO2. Lawns were resuspended in PBS, OD adjusted to ∼0.4 and heat inactivated at 57°C for 1 h and then plated to confirm bacterial heat killing. 20 μL/well of bacterial suspension was allowed to dry onto 384 well ELISA plates over 48 h in a biosafety cabinet and plates were stored at 4°C until use. Plates were blocked at room temperature for 1 h with 5% bovine serum albumin in PBS (BSA, BioShop Canada, cat. ALB001), and serum was added overnight at 4°C at a dilution of 1:4000 for cross-reactivity ELISAs or 2-fold serial dilutions for endpoint titres in 1% BSA. After washing, secondary antibody was added at 1:10,000 dilution (Jackson ImmunoResearch, cat. 115-035-003) and subsequently developed with KPL SureBlue TMB Microwell Peroxidase Substrate (SeraCare, cat. 5120-0077). Endpoint titre was calculated as the highest dilution to give a signal over background + 3 standard deviations.

### Serum bactericidal assays

Nme M982 was grown overnight on GC-Isovitalex plates at 37°C with 5% CO2. The next day, bacteria was restreaked onto fresh GC-Isovitalex plates containing 100 μmol/L deferoxamine mesylate salt (Sigma, cat. D9533) for 4 h. Bacteria were collected off the plates using a Dacron swab, resuspended in PBS++, and OD measured. Assay was set up in 40 μL total volume using ∼500 bacteria suspended in PBS++, 10% baby rabbit complement (Cedarlane, CL3441-S100) and 2-fold serial dilutions of heat inactivated terminal mouse serum. The dilution at which 50% killing was observed relative to no antibody control (rabbit complement only) in 60 minutes was reported as the SBA titre.

### hTf growth inhibition assay

*Nme* M982 was streaked onto GC-Kellogg’s plates and grown overnight at 37°C with 5% CO2. Bacteria were collected with a Dracon swab and resuspended into RPMI 1640 (Sigma, cat. R8758) to an OD600 of 0.075 and grown at 37°C with shaking at 160 rpm for 4 hours. Serum from mice immunized with their respective antigen was pooled and heat-inactivated at 56°C for 40 minutes and filtered (0.22 μm filter, Corning Costar, cat. CLS8161) to remove any aggregates and contaminants. In a 96-well round bottom plate, sera were diluted 1:20 in RPMI 1640 supplemented with a final concentration of 0.14 μM (10 μg/mL) of holo human transferrin (Sigma, cat no. T4132) to approximately an OD600 of 0.2. Bacterial growth was measured through OD600 using a Cytation 5 cell imaging multimode reader (BioTek) every 4 minutes for 10 hours at 37°C with shaking every 20 seconds.

### Western blots

Bacterial strains were grown overnight on GC-Isovitalex plates at 37 °C with 5% CO2, followed by an additional subculture on GC-Isovitalex containing 100 μM deferoxamine mesylate salt for 4 hours to induce TbpB expression. Whole cell lysates were prepared as described previously (Fegan et al., 2025). Samples were resolved on 10% SDS–PAGE and transferred to PVDF membranes. Membranes were blocked in 5% skim milk in Tris-buffered saline with 0.05% Tween (TBST) for 1 h, followed by overnight incubation at 4 °C with gentle agitation in 1:5,000 dilution of either pooled mouse or rabbit serum in 5% skim milk. Secondary antibody (either goat-anti-mouse IgG or goat anti-rabbit IgG (Jackson Immuno Research, cat. 115-035-003 or 111-035-144) was used at 1:15,000 dilution in 5% skim milk for 2 h at room temperature, after which Novex™ ECL Chemiluminescent Substrate Reagent Kit (Invitrogen) was added for band detection.

## Data availability

Coordinates of the LCL have been deposited in the Protein Data Bank under the accession code 5KKX.

## Acknowledgements

The structural analysis described in this work is based upon research conducted at the Northeastern Collaborative Access Team beamlines, which are funded by the National Institute of General Medical Sciences from the National Institutes of Health (P30 GM124165). This research used beamtime awards from the Advanced Photon Source, a U.S. Department of Energy (DOE) Office of Science User Facility operated for the DOE Office of Science by Argonne National Laboratory under Contract No. DE-AC02-06CH11357.

The authors appreciate the financial support received for the research, authorship, and/or publication of this article. This research was supported by National Institute of Health funding #R01-AI125421-01A1 and #1RO1AI141229. S.D.G. and T.F.M. are supported by the Canada Research Chair Program.

The authors would like to thank the animal support staff at the Division of Comparative Medicine at the University of Toronto for technical and welfare support for the mouse studies presented here.

## Author contributions

EAI designed, performed, analyzed murine studies, serology and cross-reactivity and SBA; JEF designed, performed, analyzed murine studies and serology and SBA; GC designed, performed and analyzed protein stability and bioinformatics studies; CC solved the crystal structure of LCL; DN performed protein purifications; NA performed growth inhibition assays; CMB and SA designed, performed and analyzed murine studies; DC performed bioinformatic analysis; LLC performed mouse studies. EAI, JEF, GC wrote the original draft; ABS, TFM, SDG provided supervision and edited the manuscript.

