## Supplemental Information for "Rationally designed minimized TbpB confers broad protection against meningococcal infection"

### Supplemental Figures

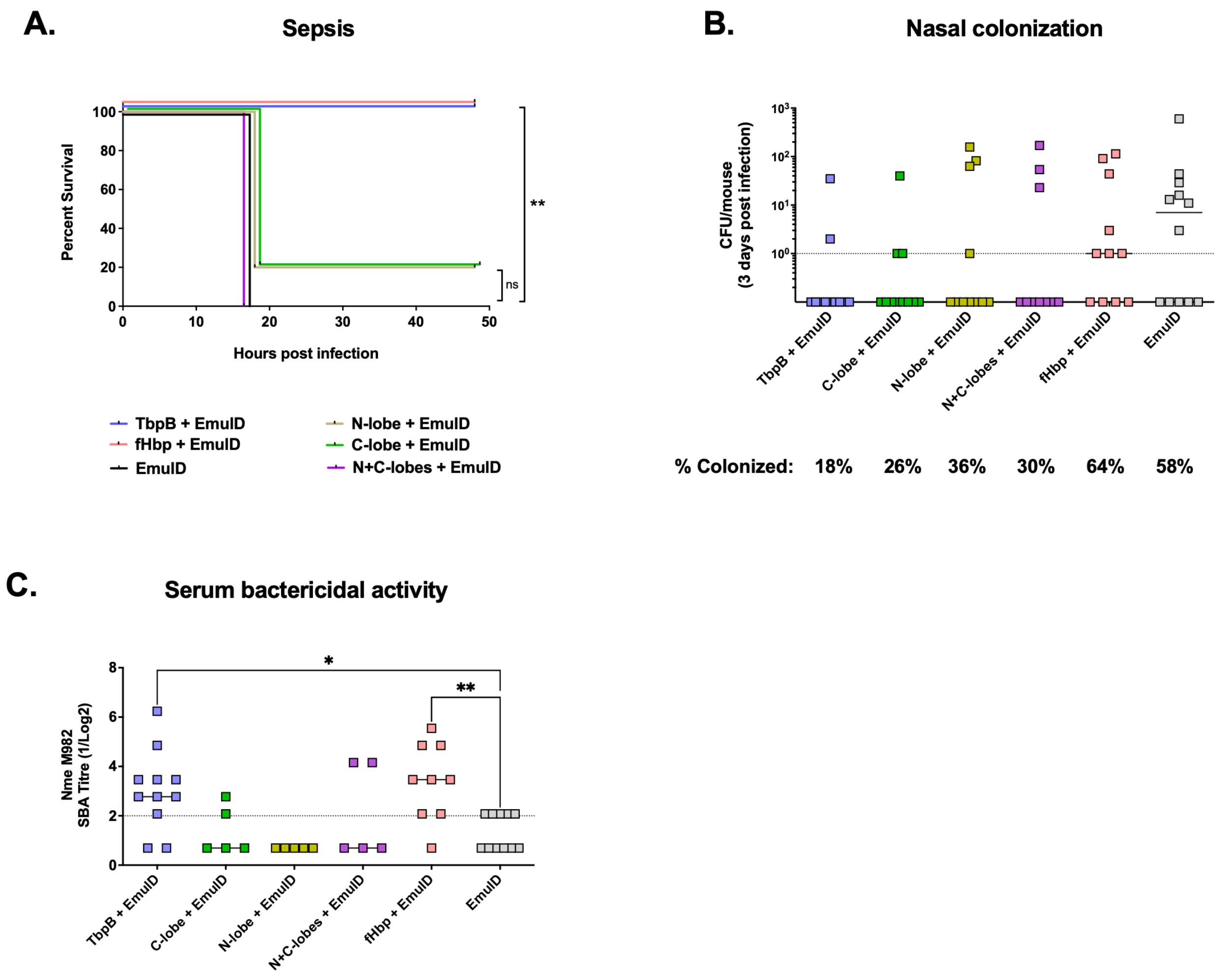


**Figure S1: Comparison of TbpB versus individual N- and C-lobes or fHbp as vaccine antigens**. A: Protection against intraperitoneal administration of a lethal dose of the homologous M982 Nme strain in C57BL/6 males immunized with TbpB, TbpB lobes, or fHbp adjuvanted with Emulsigen-D (EmulD). N=5 mice per group; p-value calculated using Log-rank (Mantel-Cox) test of each group individually against the adjuvant only control; ns, not significant; **, p<0.01. B: Graph depicting meningococcal burden in the nasopharynx of immunized hCEACAM1+/- (mCEACAM1-/-) C57BL/6 mice 3 days post nasal administration of the homologous M982 Nme strain, with the percentage of culture positive animals indicated below. C: SBA titre against iron-starved Nme M982 using terminal serum from B. N=12 for adjuvant control group, N=11 for others. B, C: Dotted lines represent lower limit of detection. Each symbol represents one animal, line at median. One-way ANOVA with Tukey's post-hoc test comparing each group to every other group performed, p<0.05 indicated. For C, log2 transformed data was used for statistical analysis. *, p<0.05; **, p<0.01.

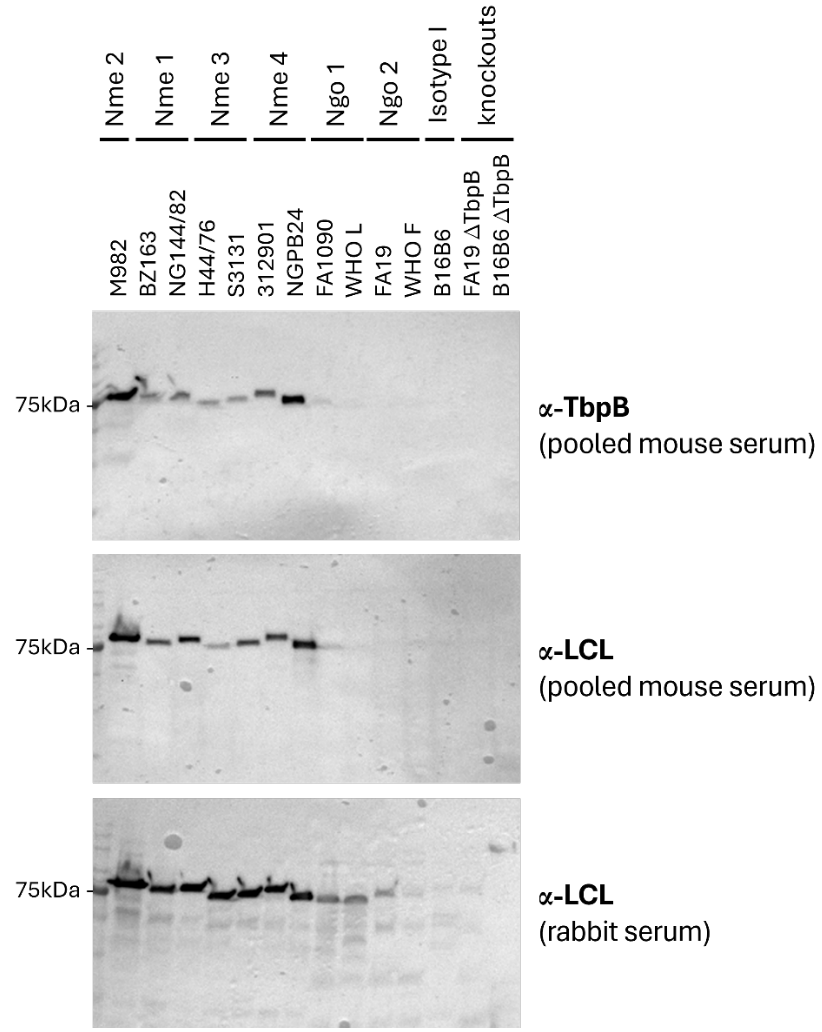


**Figure S2: Immunoblots depicting TbpB specificity.** Anti-TbpB and anti-LCL mouse serum (pooled from 8 animals per group) and anti-LCL rabbit serum (from a single animal) were used to probe whole-cell lysates prepared from iron-starved meningococcal and gonococcal strains indicated above.

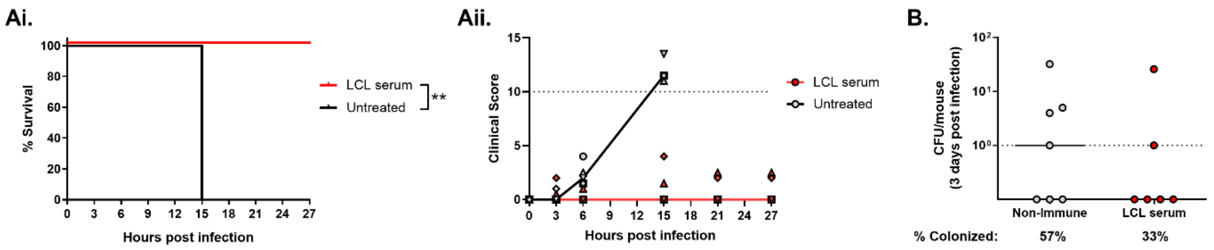


**Figure S3: Passive immunization with LCL serum.** Ai: Survival and Aii: clinical scores after intraperitoneal injection with Nme M982 in C57BL/6 mice that were untreated versus treated with inactivated rabbit LCL antiserum 6 hours prior to infection. N=5 mice per group; p-value calculated using Log-rank (Mantel-Cox) test; **, p<0.01. Dotted line at 10 depicts in the clinical score graph represents cutoff for humane endpoint. B: Meningococcal burden in the nasopharynx of hCEACAM1+/- (mCEACAM1+/+) FvB mice that were either untreated or intraperitoneally injected with LCL serum 20 hours prior to nasal infection with the homologous M982 Nme strain, with the percentage of culture positive animals indicated below. Line depicts the limit of detection.

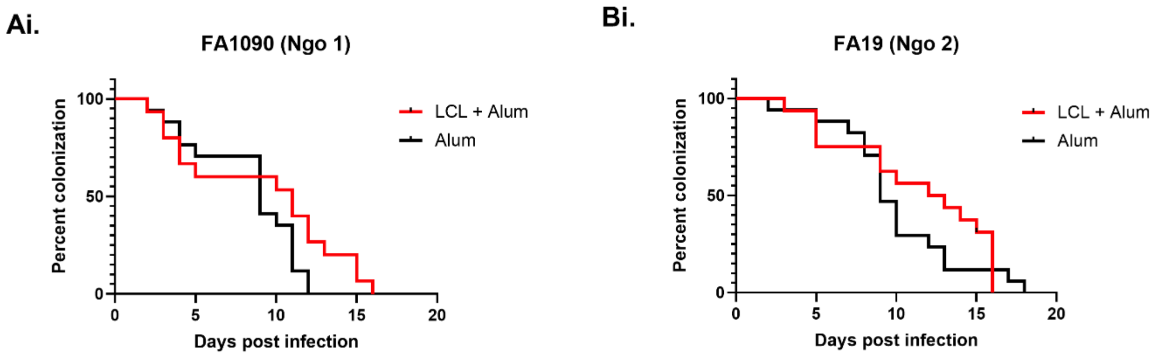


| | N | Median days colonized |
|---|---|---|
| LCL | 15 | 11 |
| Alum | 17 | 9 |

| | N | Median days colonized |
|---|---|---|
| LCL | 14 | 12.5 |
| Alum | 17 | 9 |

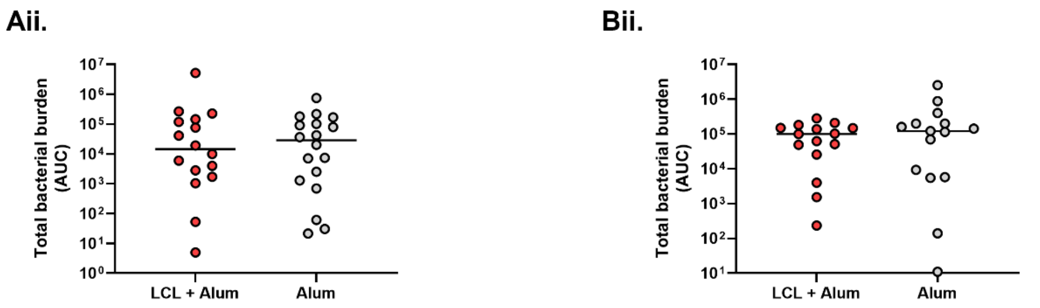


**Figure S4: Lower genital tract colonization of female mice by representative heterologous Ngo strains.** Ai, Bi: Graphs depicting % of animals that remain colonized over a three week period after vaginal inoculation with FA1090 and FA19, respectively. Groups sizes and median colonization duration indicated below. Log-rank (Mantel-Cox test) comparing Alum versus LCL curves did not yield significant p-values. Bi, Bii: Total bacterial burden of FA1090 and FA19 infected animals respectively, calculated using area under the curve (AUC) by first plotting daily CFU recovered from vaginal lavages. N=14-17 per group. Line at median. Non-parametric Mann-Whitney test did not yield significant p-values. Statistical analysis performed using GraphPad Prism 10.2.0.

**Table S1: Refinement statistics for the LCL crystal structure.**

| | **LCL** |
|---|---|
| PDB Code | 5kkx |
| Phasing Method | Molecular replacement |
| Data Collection | |
| Wavelength | 0.9795 |
| Resolution range | 33.23 - 1.899 (1.95 - 1.9) |
| Space group | P 4 21 2 |
| Cell Dimensions | |
| a, b, c (Å) | 132.939 132.939 64.353 |
| α, β, γ (°) | 90 90 90 |
| Unique reflections | 45698 (3162) |
| Completeness (%) | 99.37 (98.20) |
| Wilson B-factor | 34.56 |
| Refinement | |
| Resolution | 2.1 |
| Reflections used in refinement | 45698 (3162) |
| Reflections used for R-free | 1995 (138) |
| R-work | 0.1825 (0.3598) |
| R-free | 0.2166 (0.3954) |
| Number of non-hydrogen atoms | 3842 |
| macromolecules | 3638 |
| ligands | 12 |
| solvent | 192 |
| Protein residues | 462 |
| R.m.s. deviations | |
| RMS(bonds) | 0.007 |
| RMS(angles) | 1.1 |
| Ramachandran favored (%) | 95.35 |
| Ramachandran allowed (%) | 4.65 |
| Ramachandran outliers (%) | 0 |
| Rotamer outliers (%) | 1.82 |
| Clashscore | 8.45 |
| Average B-factor | 52.34 |
| macromolecules | 52.36 |
| ligands | 68.4 |
| solvent | 50.94 |
